# Interleukin-1α and Leukemia Inhibitory Factor Promote Extramedullary Hematopoiesis

**DOI:** 10.1101/2022.07.07.499256

**Authors:** Derek A.G. Barisas, Minseo Kim, Madhav Subramanian, Ashraf Ul Kabir, Karen Krchma, Jun Wu, Bernd H. Zinselmeyer, Colin L Stewart, Kyunghee Choi

## Abstract

Extramedullary hematopoiesis (EMH) expands hematopoietic capacity outside of the bone marrow in response to inflammatory conditions, including infections and cancer. Because of its inducible nature, EMH offers a unique opportunity to study the interaction between hematopoietic stem and progenitor cells (HSPCs) and their niche. In cancer patients, the spleen frequently serves as an EMH organ and provides myeloid cells that may worsen pathology. Here, we examined the relationship between HSPCs and their splenic niche in EMH secondary to solid tumor. We identify an inflammatory gene signature characterized by TNFα expression in HSPCs. We show a role for IL-1α in producing this gene signature and TNFα expression in HSPCs in activating splenic niche activity. We also demonstrate that tumor-derived Leukemia Inhibitory Factor (LIF) induces proliferation of splenic niche cells. IL-1α and LIF display cooperative effects in activating EMH and are both upregulated in some human cancers. Together, these data expand avenues for developing niche-directed therapies and further exploring EMH accompanying inflammatory pathologies like cancer.

## Introduction

Hematopoiesis produces differentiated cell types of the blood and immune systems from hematopoietic stem and progenitor cells (HSPCs). Organismal changes such as disease can modulate the location and cellular output of hematopoiesis [1, 2].

Expansion of hematopoiesis outside of the bone marrow (BM), known as extramedullary hematopoiesis (EMH), accompanies pathologic states and occurs mainly within the spleen and liver. Long underappreciated in human disease, EMH is now beginning to be recognized as important component to multiple hematologic and non-hematologic disease [3, 4]. The induction of EMH requires mobilization of HSPCs from the BM by cytokines, such as ligands for CXCR2 including CXCL1 and CXCL2 [5–7]. Clinically, EMH presents in a diverse set of solid tumors, including breast, lung, renal, colon, gastric, pancreatic, and prostate cancer [8, 9]. Of particular interest is EMH in the spleen due to the organ’s role in supplying myeloid cells during multiple injury and disease states and its frequent involvement in cancer patients [8, 10, 11].

Myeloid-biased differentiation is a response of hematopoiesis to inflammatory signals, including IL-1β, TNFα, and G-CSF [12–14]. Enhanced myelopoiesis, characteristic of EMH, can exacerbate diseases like solid tumors, arthritis, and myocardial infarction by increasing the number of cells that drive pathology [9, 15, 16]. Clinically, increased myeloid cell production can be measured by an increased ratio of neutrophils (PMNs) to lymphocytes in the peripheral blood (PB). Across multiple tumor types, including breast, colon, pancreatic, and gastric cancer, as well as a systematic review of all cancer types, a high neutrophil-to-lymphocyte ratio in the peripheral blood is a poor prognostic factor for survival [17–20].

Like other stem cells, hematopoietic stem cells rely on supporting cell types known as the niche [21, 22]. Essential to hematopoietic niche function is the production of membrane-bound KIT ligand, a key growth factor for HSPCs [23–25]. Additionally, the niche must produce factors to attract and adhere HSPCs. CXCL12 is a critical chemotactic factor for HSPCs within the BM niche while VCAM-1 is central to adherence through interactions with VLA-4 and other integrins on HSPCs [26, 27].

Among hematopoietic niche cell types, perivascular stromal cells play a central role through their production of KIT ligand and CXCL12 [28, 29]. Mesenchymal stem cells have been shown to exert niche function in both mice and humans [30, 31]. Despite their importance to the niche, demarcating cells as perivascular stromal cells has been tricky. However, several schemas have recognized PDGFRα as an important marker and noted their co-expression of PDGFRβ [32–34]. This PDGFRα+/β+ surface phenotype matches mesenchymal stem cells as identified by single-cell RNA-sequencing of limb muscles [35, 36].

Significant advances have been made in delineating the BM niche and HSPC interaction at homeostasis [21, 22]. However, although perivascular stromal cells have been appreciated as contributing to the splenic niche at homeostasis [37, 38], HSPC niches outside of the BM that support EMH are less well-understood. Here we demonstrate the importance of splenic EMH in producing PMNs during a mouse model of breast cancer and identify a novel inflammatory phenotype for HSPCs conducting EMH. We delineate cytokine communication between IL-1α-inflamed, splenic HSPCs expressing TNFα and their splenic niche. We also investigate a parallel mode of cytokine communication between tumor cells and the splenic niche through Leukemia Inhibitory Factor (LIF). Both pathways may increase the myelopoietic capacity of the spleen during inflammatory pathologies such as solid tumors.

## Results

### Murine cancer models have expanded splenic hematopoiesis, often with bias towards myelopoiesis

The MMTV-PyMT mouse is a genetic model of breast cancer where tumors develop *in situ* due to an oncogene under the control of a promoter expressed primarily in mammary epithelium [39]. Mice with tumors experience neutrophilia and a drastic increase in spleen weight, spleen cellularity, and splenic HSPCs as measured by c-Kit^+^/Sca-1^+^/Lineage^-^ (KSL) and granulocyte-monocyte precursor (GMP) amounts (Fig. S1A-E). These changes occur with minimal effects to the splenic common lymphoid progenitors (CLP) (Fig. S1F) or the BM compartment (Fig. S1G-I). To provide more experimental control than the genetic model, we developed a heterotopic tumor model using a PyMT-B6 cell line derived from tumors of a B6/J syngeneic MMTV-PyMT mouse [40]. PyMT-B6 tumors in female mice aged 8 to 16 weeks, a gender and age range used in further experimentation unless otherwise stated, produced neutrophilia in animals 21 days post-injection (Fig. 1A). Similarly, these mice have increased spleen weight, spleen cellularity, and HSPC (KSL) and GMP amounts, in total and as a percent of CD45^+^ cells (Fig. 1B-G). This effect was not mirrored in the BM compartment or the splenic common lymphoid progenitors (Fig. 1H-K). Additionally, increased CFU-GEMM colony numbers were identified in the PB of PyMT-B6 bearing animals compared to controls, suggesting mobilization of HSPCs outside of the BM (Fig. 1L). Together, data from both a genetic and transplantation model identify the spleen as a site of profound HSPC expansion coincident with increased granulocytes and primitive hematopoietic progenitors in the PB.

**Figure 1.**
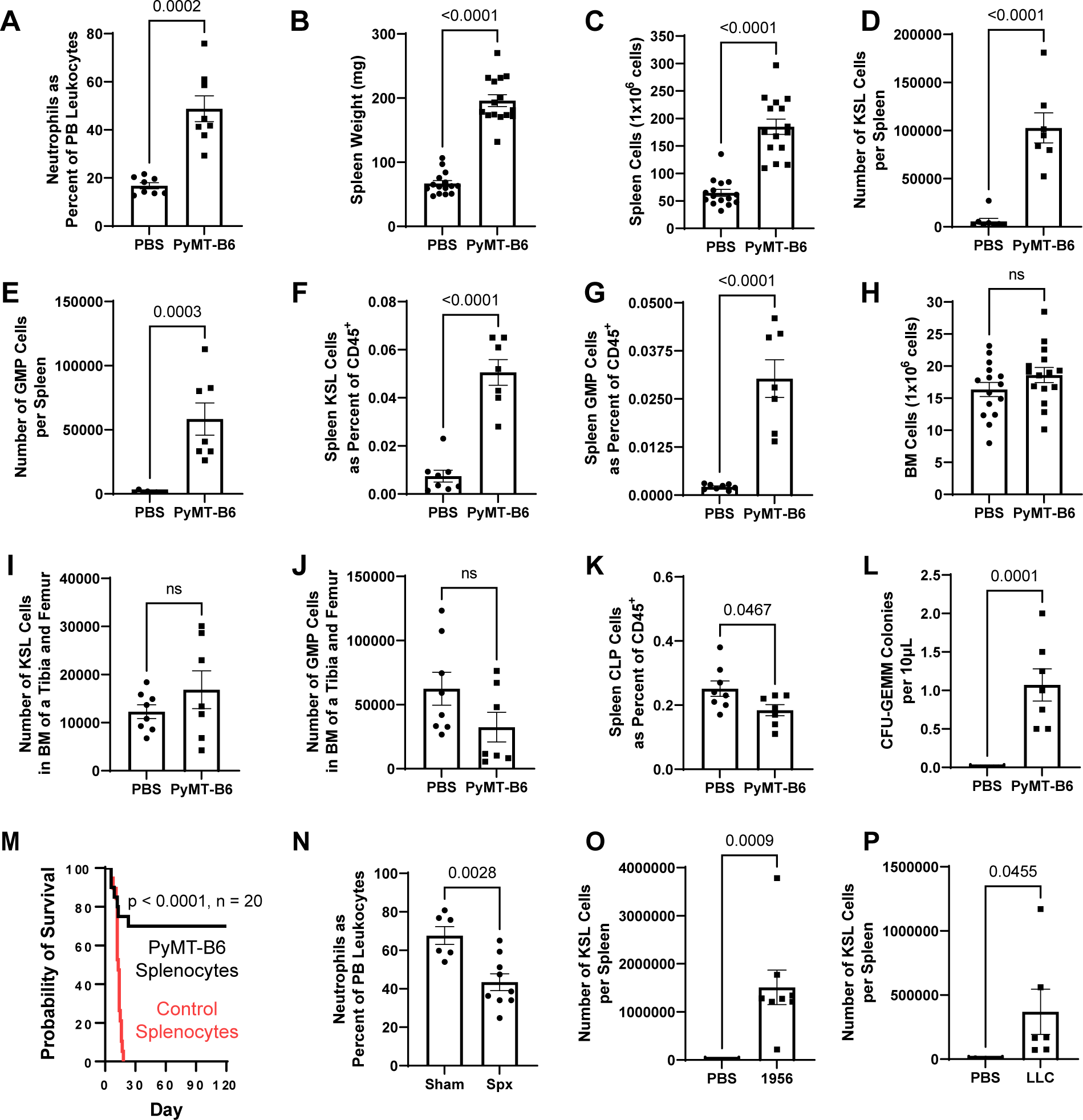
Expansion of splenic hematopoiesis is required for neutrophilia in PyMT-B6. (A - K) 21 days after injection of 5 x 10^5^ PyMT-B6 tumor cells injected subcutaneously compared to control animals injected with phosphate buffered saline, PMNs in the PB as a percent of total leukocytes (A, n = 8), splenic weight (B, n = 15), splenic cellularity (C, n = 15), KSL cells per spleen (D, n = 7-8), GMP cells per spleen (E, n = 7-8), KSL cells as a fraction of total splenic CD45^+^ cells (F, n = 7-8), GMP cells as a fraction of total splenic CD45^+^ cells (G, n = 7-8), BM cellularity per leg (H, n = 15), KSL cells per BM of a leg (I, n = 7-8), BM GMP cells per BM of a leg (J, n = 7-8), CLP cells as a fraction of total splenic CD45^+^ cells (K, n = 7-8). (L) CFU-GEMM colonies within 10μL of PB, 21 days of PyMT-B6 tumor compared to PBS injected controls (n = 7-8). (M) Survival of 9.5 Gy irradiated mice receiving splenocytes from mice with 21 days of PyMT-B6 tumor or control mice injected with PBS (n = 20, significance assigned by Mantel-Cox test). (N) PMNs in the PB as a percent of total leukocytes with 21 days of PyMT-B6 tumor following splenectomy or sham surgery (n = 7-8). (O) KSL cells per spleen 17 days after injection of 2 x 10^6^ 1956 tumor cells injected subcutaneously compared to control animals injected with phosphate buffered saline (n = 8). (P) KSL cells per spleen 16 days after injection of 5 x 10^5^ LLC tumor cells injected subcutaneously compared to control animals injected with phosphate buffered saline (n = 6-7).

Enhanced survival of irradiated CD45.2 mice transplanted with CD45.1 splenocytes of PyMT-B6 bearing animals compared to mice receiving control splenocytes validated the stem cell function of these splenic HSPCs (Fig. 1M). Furthermore, these animals had

>99% of donor-derived myeloid and lymphoid lineages in the PB when analyzed at 4 months of transplantation. Splenectomized animals had reduced PMN percentages in their PB after 21 days of PyMT-B6 tumor compared to sham surgery controls (Fig. 1N). To generalize our findings about expanded splenic hematopoiesis to other cancer models, heterotopic models of 1956 sarcoma and LLC lung carcinoma were investigated and found to significantly expand HSPC and GMP populations with a variable response to peripheral neutrophilia (Fig. 1O, P; SFig. 1J-M) [41–43]. Together, these data indicate that breast cancer induces expansion of splenic hematopoiesis that is necessary for neutrophilia and this expanded capacity is generalizable to other murine cancer models.

### HSPCs conducting EMH express an inflammatory gene profile

To characterize the transcriptional and cell composition changes induced by the PyMT- B6 tumor, we performed single-cell RNA-sequencing (scRNA-seq) of splenic and BM cells enriched for HSPCs in mice with or without PyMT-B6 tumor. The Lin^-^/c-Kit^+^/Sca- 1^+^/CD34^+^ cells representing the hematopoietic stem and progenitor cells (Fig. S2A-I, Fig. 2A-C), while rarely present in the control spleen, were well-represented in the spleen of the tumor-bearing mice (Fig. 2D). The Lin^-^/c-Kit^+^/Sca-1^+^/CD34^+^ cells from the spleen of tumor-bearing animals expressed a unique gene signature, including *Tnf*, *Cxcl2*, *Nfkbiz*, *Nfkbia,* compared to control BM (CBM) cells or tumor-bearing BM (TBM) cells (Fig. 2E-H). We chose to focus on the comparison between control bone marrow and HPSCs associated with EMH in the spleen to address questions about the functional differences between homeostatic HSPCs and those associated with pathology and residing in extramedullary sites. The up-regulation of these four genes in tumor-bearing, spleen (TS) Lin^-^/CD34^+^ HSPC compared to the same population in the homeostatic BM was confirmed by RT-qPCR (Fig. 2I-L). Increased TNFα protein expression was identified within the HSPC fraction of TS relative to CBM by flow cytometry (Fig. 2M-O). Together, these data demonstrate that PyMT-B6 tumor presence activates an inflammatory gene signature within splenic HSPCs that induces TNFα expression by these cells.

**Figure 2.**
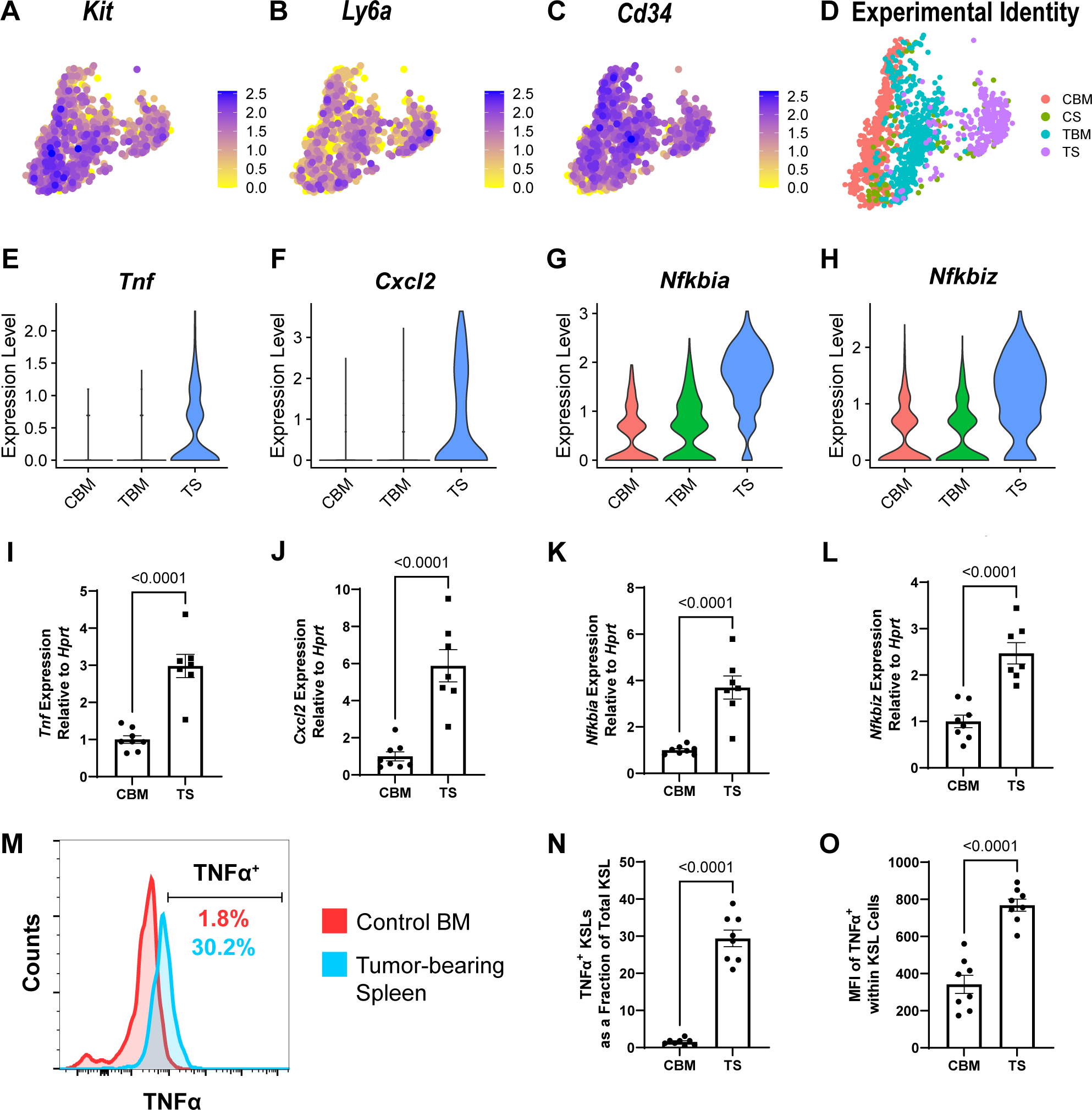
HSPCs from tumor-bearing mice display an inflammatory gene signature. (A – D) UMAP projection of the HSPC population in scRNA-seq data colored by expression of *Kit* (A), *Ly6a* (B), *Cd34* (C) and by experimental origin (D, CBM – Control BM, CS – Control Spleen, TBM – Tumor-bearing BM, TS – Tumor-bearing spleen). (E – H) Violin plot of expression of *Tnf* (E), *Cxcl2* (F), *Nfkbia* (G), and *Nfkbiz* (H) in the HSPC population in scRNA-seq data from CBM, TBM, or TS. (I - L) RT-qPCR expression data of *Tnf* (I), *Cxcl2* (J), *Nfkbia* (K), *Nfkbiz* (L) from Lin^-^/Flk1^-^/CD34^+^ cells from CBM or TS. (M) Representative histogram of TNFα expression in KSL cells from CBM or TS. (N) Fraction of KSL cells from CBM or TS that are TNFα^+^ (n = 8). (O) Mean fluorescent intensity of TNFα staining in KSL cells from CBM and TS (n = 8).

### PyMT-B6-produced IL-1α acts on HSPCs to express TNFα

TNFα expression in splenic HSPCs of PyMT-B6 bearing mice hints at the presence of tumor-derived upstream mediators. One often reported cytokine subfamily upstream of TNFα is IL-1α and IL-1β [44]; of which, IL-1α, but not IL-1β, is produced during MMTV-PyMT tumor pathology [45]. We confirmed that mice bearing PyMT-B6 tumors have elevated circulating levels of IL-1α (Fig. 3A) and that PyMT-B6 cells release IL-1α *in vitro* (Fig. 3B). Correspondingly, IL-1 receptor was identified as being constitutively expressed in HSPCs by scRNA-seq and RT-qPCR, although its expression was somewhat lower in tumor-bearing splenic HSPCs compared to control BM HSPCs (Fig. 3C-D). Injection of IL-1α into mice was sufficient to induce neutrophilia, increase splenic HSPC fraction, and increase TNFα expression in splenic HSPCs compared to control BM HSPCs (Fig. 3E-G). Reciprocally, deletion of *Il1a* from PyMT-B6 cells led to decreased expression of TNFα in HSPCs and decreased total splenic GMP cells compared to the parental line (Fig. 3H, I, Fig. S2J, K). The parental line includes deletion of G-CSF, a cytokine shown to produce myeloid-biased hematopoiesis and EMH that may obscure the effects of other cytokines [46]. Together, these data indicate that PyMT-B6 IL-1α induces a novel inflammatory phenotype in HSPCs associated with tumor-induced EMH.

**Figure 3.**
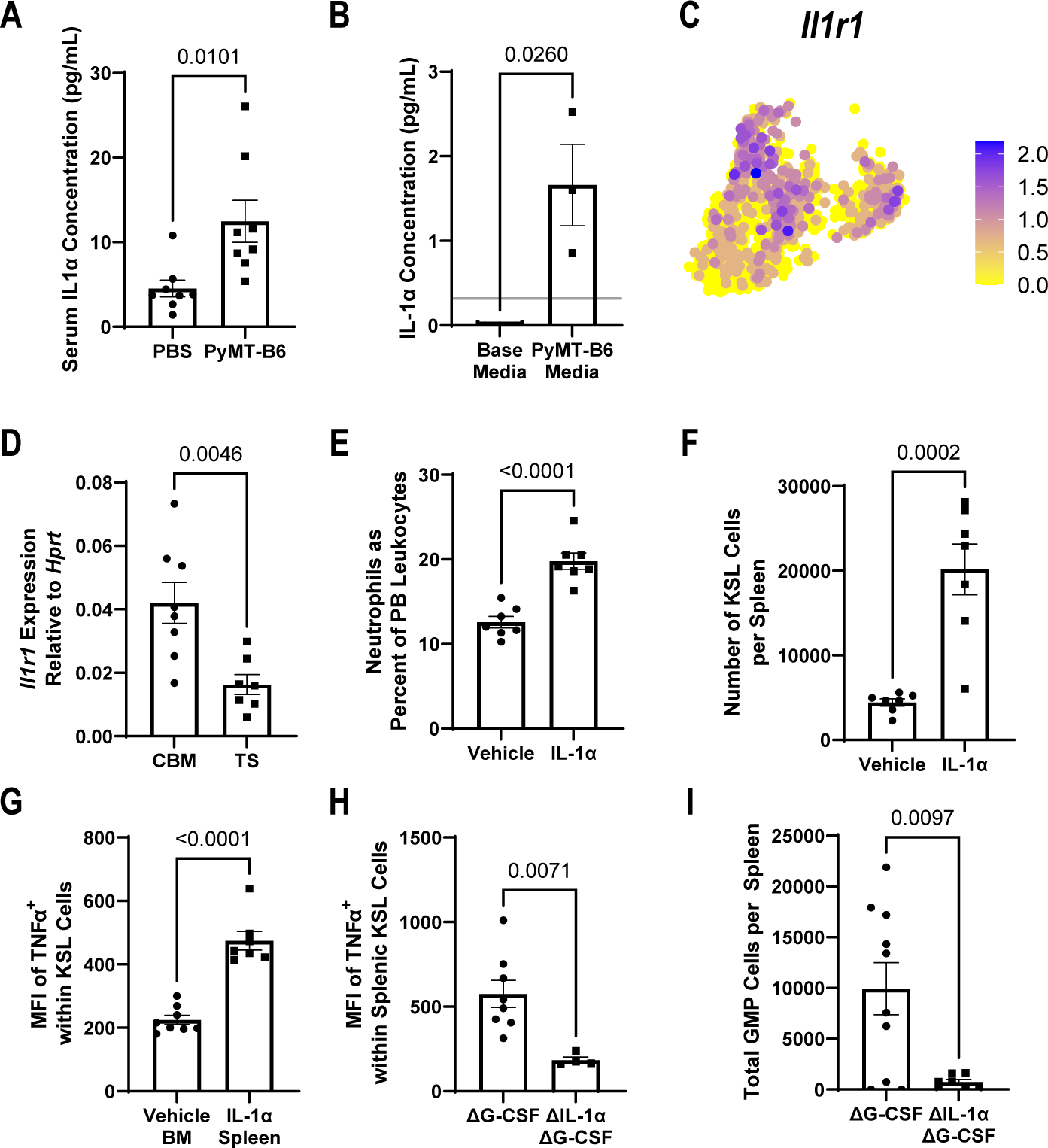
Tumor-derived IL-1α activates TNFα production in splenic HSPCs. (A) IL- 1α concentration from serum of mice with or without 21 days of PyMT-B6 tumor (n = 8). (B) IL-1α concentration from PyMT-B6 base or conditioned media (n = 3). (C) UMAP projection of HSPCs in scRNA-seq data colored by *Il1r1* expression. (D) RT-qPCR expression data of *Il1r1* from Lin^-^/Flk1^-^/CD34^+^ cells from control bone marrow (CBM) or PyMT-B6 tumor bearing spleen (TS). (E – F) In mice 24 hours after intravenous (i.v.) injection of 500ng IL-1α or vehicle, PMNs in the PB as a percent of total leukocytes in mice (E, n = 7), KSL cells per spleen, (F, n = 7). (G) Average mean fluorescent intensity of TNFα staining in KSL cells from vehicle injected BM or 500ng IL-1α injected spleen. (n = 7-8). (H – I) 28 days after subcutaneous injection of 2.5 x 10^5^ PyMT-B6 ΔG-CSF parental cells or ΔG-CSF ΔIL-1α cells, Average mean fluorescent intensity of TNFα staining in KSL cells (H, n = 7-13), GMP cells per spleen (I, n = 7-13).

### TNFα induces EMH through splenic niche cells

Due to the concurrent TNFα production by splenic HSPCs and splenic EMH accompanying PyMT-B6 tumors, we assessed whether TNFα from HSPCs could induce EMH by activating local niche cells. Administration of a single dose of TNFα was sufficient to increase HSPC and GMP fractions in the spleen within 24 hours and to produce neutrophilia (Fig. 4A-C). To test whether niche cells respond to TNFα, we needed to identify potential niche cells within the spleen. Reanalysis of a BM niche cell scRNA-seq dataset [47] identified *Pdgfra+/Pdgfrb+* stromal (ABS) cells as being the most strongly KIT ligand positive cell population and expressing a TNFα receptor (Fig. 4D-G, cluster 3, S3A-I). Using a novel method (see Materials and Methods section), we cultured ABS cells from the spleen and validated their expression of membrane KIT ligand by flow cytometry (Fig. 4H, I). To investigate the niche functionality of these ABS cells, 5,000 live BM Lin^-^/c-Kit^+^ (KL) cells, of which around 20% were also Sca-1^+^, were sorted into 24-wells with or without confluent ABS cell cultures. After 7 days of co-culture, a large population of small, spherical cells grew on top of the ABS monolayer (Fig. S3J, K). Upon flow cytometric evaluation, these co-cultures contained a population of CD45^+^/Lin^-^/c-Kit^+^/Sca-1^+^ cells (Fig. S3I). To test whether the hematopoietic component of these co-cultures maintained stem cell capacity, first, CD45.1 KL cells were sorted into plates and cultured for 7 days with or without ABS stromal before transplanting them into irradiated CD45.2 recipient mice. Compared to mice receiving KL cells cultured without ABS cells, mice that received KL cells cultured on ABS cells had significantly improved survival, indicating the maintenance of repopulating units *in vitro* (Fig. 4J). Analysis of the peripheral blood from surviving transplant mice indicated donor derived hematopoietic cells constituted more than 90% of all CD45^+^ cells after one month. Second, colony forming unit activity was compared between KL cells grown with or without ABS cells for 7 days. After 7 days, more primitive precursor activity, as measured by CFU-GEMM colony formation, was nearly absent from cells without co-culture but preserved in cells grown in co-culture (Fig. 4K). Additionally, CFU-GEMM colonies were observed until at least 21 days in co-culture. Having established genuine HSPC niche activity, we wanted to understand how ABS cells might change phenotypically in response to HSPC cytokines. Following TNFα addition to culture medium, splenic ABS cells increased HSPC-adherent VCAM-1 expression and released the HSPC active chemokine CXCL1 while maintaining baseline CXCL12 release (Fig. 4L-N). Together, these data suggest that TNFα produced by HSPCs during tumor presence can act locally on ABS niche cells to increase the capacity of the splenic niche to support hematopoiesis.

**Figure 4.**
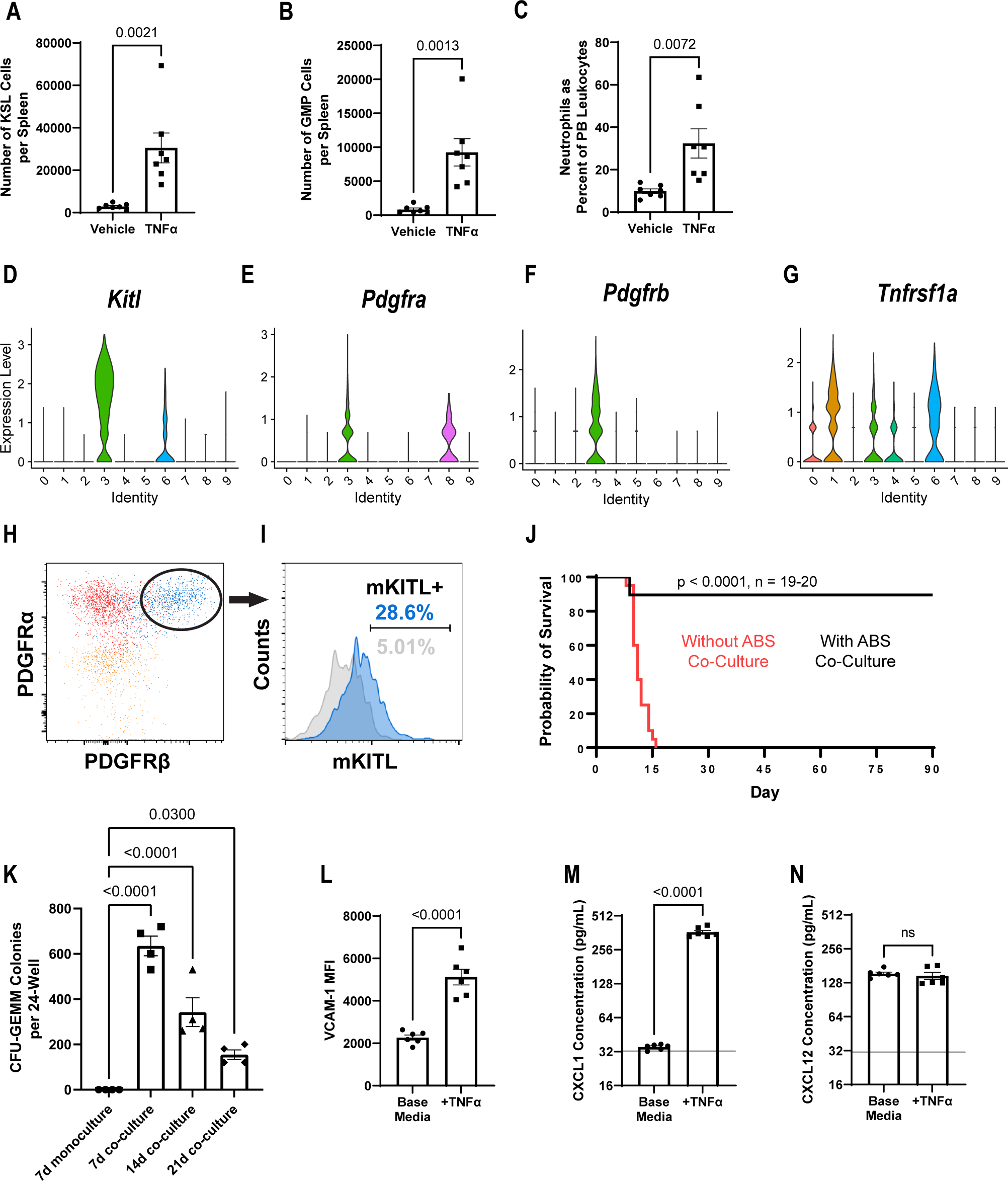
TNFα activates stromal cells in the spleen to induce EMH. (A – C) In mice 24 hours after i.v. injection of 2μg TNFα or vehicle, KSL cells per spleen (A, n = 7), GMP cells per spleen (B, n = 7), PMNs in the PB as a percent of total leukocytes (C, n = 7). (D – G) Violin plot of expression of *Kitl* (D), *Pdgfra* (E), *Pdgfrb* (F), *Tnfrsf1a* (G) in reanalyzed scRNA-seq data from Tikhonova *et al.* 2019 of bone marrow niche cell types (0 – HSC, 1 – endothelium (EC), 2 – Proliferating CD45^+^, 3 – ABS cell, 4 – GMP, 5 – CLP, 6 – Sca-1^+^ EC, 7 – B-cell progenitor, 8 – Osteoblast, 9 – RBC Progenitor). (H) Representative dot plot comparing splenic ABS cells stained solely for viability, a PDGFRβ fluorescence minus one sample, and a fully stained sample. (I) Representative histogram of membrane KITL expression in splenic ABS cells. (J) Survival of 9.5 Gy irradiated mice receiving the cell products of c-Kit^+^Lin^-^ cells grown with or without ABS cell co-culture for 7 days (n = 19-20, significance assigned by Mantel-Cox test). (K) Representative quantification of CFU-GEMM colonies per 24-well of the cell products of c-Kit^+^Lin^-^ cells grown with or without ABS cell co-culture for 7 days and with ABS cell co-culture for 14 and 21 days. (n = 4 replicates per group, significance assigned by one-way ANOVA with multiple comparison tests against the 7d monoculture group). (L – N) In splenic ABS cells treated for 24 hours with or without 2.5ng/mL TNFα, representative mean fluorescent intensity of VCAM-1 (L, n = 6), representative CXCL1 concentration (M, n = 6), representative CXCL12 concentration (N, n = 6).

### Tumor-derived Leukemia Inhibitory Factor activates splenic EMH

Given the indirect interaction between tumor cells and splenic niche cells through inflamed HSPCs, we were interested in the potential of a direct interaction between tumor and splenic niche cells. Preliminary analysis of a 44-member cytokine array on serum from MMTV-PyMT tumor-bearing animal compared to littermates identified Leukemia Inhibitory Factor (LIF), an IL-6 family member, as being significantly and consistently upregulated by the presence of tumors (Fig. S4A). The presence of LIF in the serum of PyMT-B6 bearing animals and the production of LIF by PyMT-B6 cells in culture was independently confirmed (Fig. 5A, B). Previous work has identified LIF as having an active role in promoting and maintaining hematopoiesis in the spleen [48, 49]. We tested whether LIF might have a role in cancer-induced EMH by generating a lentiviral expression vector for murine LIF and injecting mice intravenously to induce systemic LIF overexpression. Compared to empty lentiviral vectors, LIF overexpression induced neutrophilia and a robust expansion of HSPC and GMP cells in the spleen (Fig. 5C-E). Correspondingly, deletion of *Lif* from Δ*Csf3* PyMT-B6 cells lead to decreased levels of splenic HSPCs and GMPs compared to the Δ*Csf3* parental line (Fig. 5F, Fig. S4B-C). These data identify LIF as a tumor-secreted factor which is sufficient to induce myeloid-biased expansion of hematopoiesis within the spleen.

**Figure 5.**
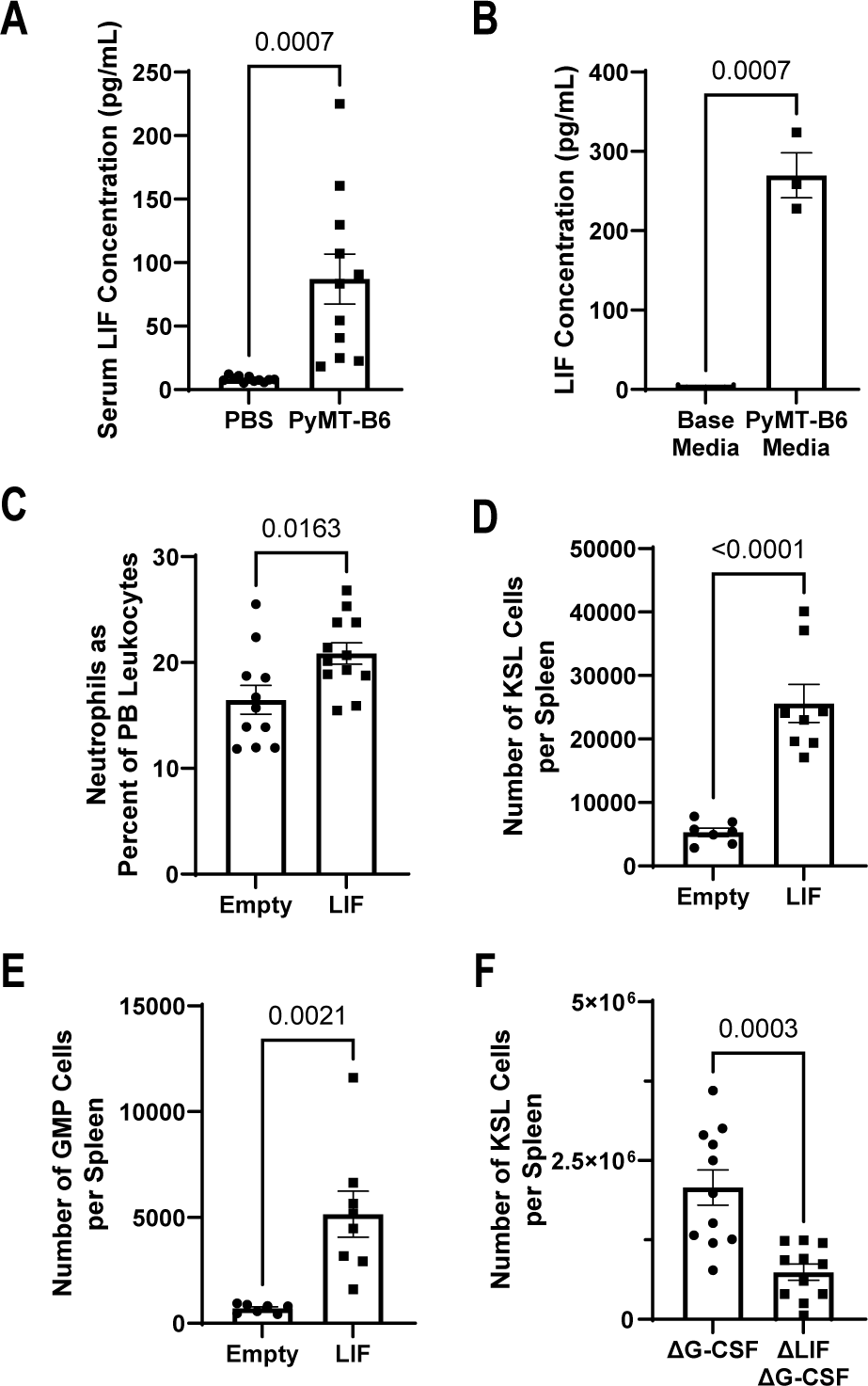
Tumor-produced Leukemia Inhibitory Factor induces EMH. (A) LIF concentration from serum of mice with or without 21 days of PyMT-B6 tumor (n = 11). (B) LIF concentration from base media or PyMT-B6 conditioned media (n = 3). (C – E) In mice with 10 days of LIF overexpression or empty vector control, fraction of PMNs in the PB as a percent of total leukocytes (C, n = 11-12), KSL cells per spleen (D, n = 7-8), GMP cells per spleen (E, n = 7-8) (F) 28 days after subcutaneous injection of 2.5 x 10^5^ PyMT-B6 ΔG-CSF parental cells or ΔG-CSF ΔLIF cells, KSL cells per spleen (F, n = 12).

### Leukemia Inhibitory Factor induces splenic stromal niche cell proliferation

Having identified the capability of LIF to expand splenic hematopoietic capacity, we sought to define a cellular mechanism for its effect. Re-examination of niche scRNA-seq data [47] identified both *Kitl* expressing clusters, *Cdh5+/Ly6a*+ endothelial and ABS cells, as expressing LIF receptor (LIFR) (Fig. 6A, cluster 6 vs 3, respectively). By inducing LIF overexpression by lentivirus in Cdh5-Cre^+^/Lifr^fl/fl^ mice and littermate controls, we could exclude endothelial cell contribution to LIF-induced EMH (Fig. 6B).

**Figure 6.**
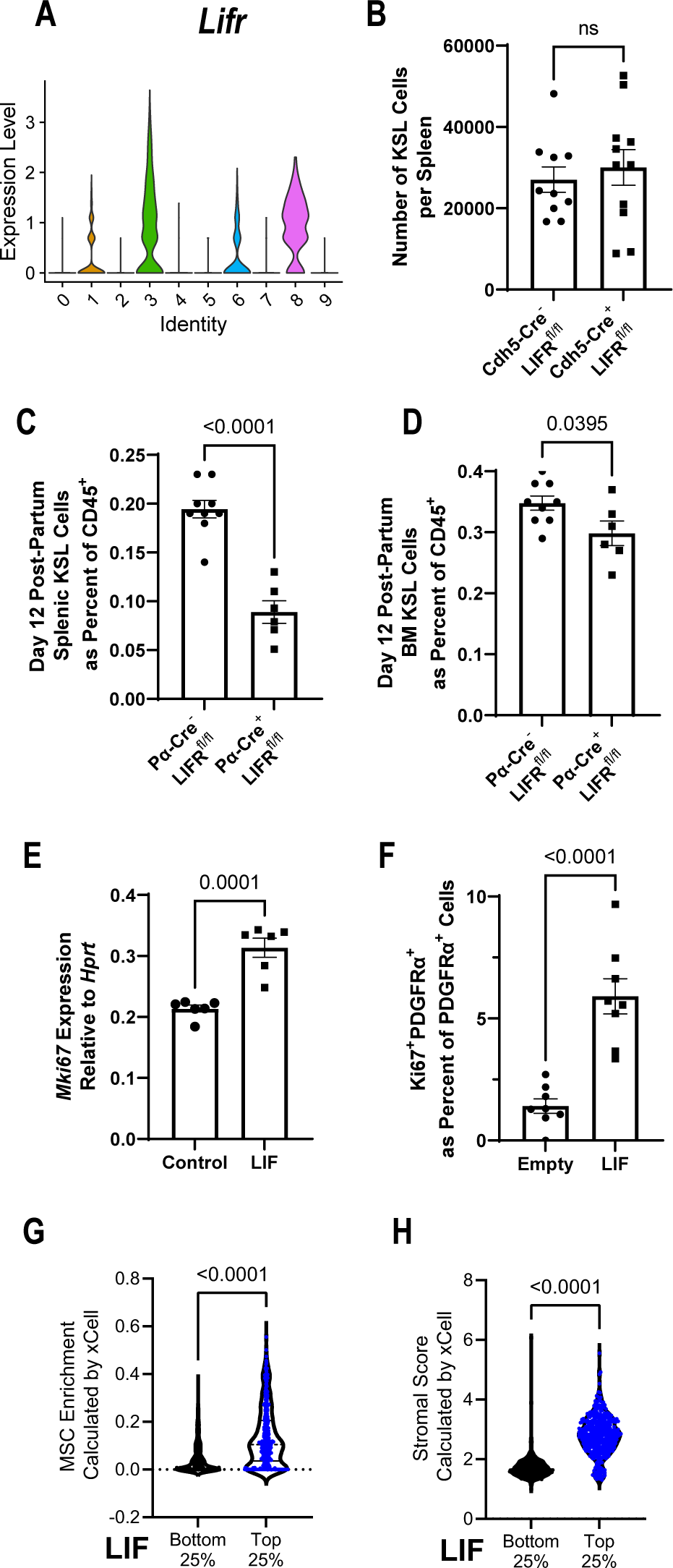
LIF directly expands the splenic niche. (A) Violin plot of expression of *Lifr* in reanalyzed scRNA-seq data from Tikhonova *et al.* 2019 of bone marrow niche cell types (0 – HSC, 1 – endothelium (EC), 2 – Proliferating CD45^+^, 3 – ABS cell, 4 – GMP, 5 – CLP, 6 – Sca-1^+^ EC, 7 – B-cell progenitor, 8 – Osteoblast, 9 – RBC Progenitor). (B) In LIFR^flox^ mice with LIF overexpression and Cdh-Cre^+^ or Cdh5-Cre^-^, KSL cells per spleen (n = 10-11, contains male mice). (C – D) In day 12 post-partum LIFR^flox^ mice with PDGFRα-Cre^+^ or PDGFRα-Cre^-^ littermates, KSL cells as a fraction of total splenic CD45^+^ cells (C, n = 6-9, contains male mice), KSL cells as a fraction of total bone marrow CD45^+^ cells (D, n = 6-9, contains male mice). (E) Representative RT-qPCR expression data of *Mki67* from splenic ABS cells treated for 72 hours with 20ng/mL LIF (n = 6). (F) Fraction of splenic PDGFRα+ cell that are Ki67+ by immunofluorescence with 7 days of LIF overexpression or empty vector lentivirus control (n = 8). (G – H) Enrichment of MSCs (G) and stromal scoring (H) as calculated by xCell from RNA-seq data of human tumors split by top and bottom quartile of LIF expression (n = 416-417).

We generated mice with LIFR deletion within the PDGFRα^+^ population to assess the involvement of splenic ABS cells in LIF response. Pdgfra-Cre^+^/Lifr^fl/fl^ mice were born at expected frequencies but died before weaning due to a failure to thrive, a similar but less severe phenotype than the constitutive knock out mouse (Fig. S4D, E) [50]. Despite the lethality at around the weaning, conditional knockouts are still alive at days 12 post- partum, a time point when the spleen still shows active hematopoiesis [51]. We found that Pdgfra-Cre^+^/Lifr^fl/fl^ mice had reduced HSPCs specifically within the spleen compared to the bone marrow and littermate controls at this time (Fig. 6C-D). This suggests that the LIF-LIFR axis in PDGFRα^+^ cells is indispensable for maintenance of hematopoiesis specifically within the spleen, and therefore we were interested in potential mechanisms. Previous studies showed that LIF induces proliferation of PDGFRα+ oligodendrocyte precursor cells and osteoblast precursors [52, 53]. We added LIF to splenic ABS cultures and found increased markers of proliferation (Fig. 6E). To confirm this finding *in vivo*, we quantified the fraction of Ki67^+^ nuclei of PDGFRα^+^ cells in the spleen with or without lentiviral LIF overexpression using immunofluorescence and found an increase in Ki67^+^ PDGFRα^+^ cells with LIF overexpression (Fig. 6F, Fig. S4F). Additionally, we found the close association of PDGFRα^+^ cells with Kit^+^ progenitors in the spleen after LIF overexpression by confocal imaging (Fig. S4G).

Our data suggest that LIF expression expands distal stromal components in mouse models. To investigate whether LIF expression in human cancer correlates with local stromal populations, we analyzed RNA-sequencing data from a collection of sources [54]. Consistent with our mouse data, tumors in the highest quartile of LIF expression had significantly higher amounts of MSCs, fibroblasts, and stromal scores compared to the lowest quartile, with only a modest increase in the endothelial fraction between the two groups (Fig. 6G-H, S4H-I). Together, these data suggest that ABS cells form an expandable niche in the spleen in direct response to tumor-derived LIF and that this cancer-stromal interaction may operate in human tumors as well.

### IL-1α and LIF have a cooperative myelopoietic response in mice and are co-expressed in human cancers

Due to their independent mechanisms in activating the splenic niche, we determined if the interaction of IL-1α and LIF would increase myelopoietic output. To this end, we first injected mice with lentiviral constructs that were either empty or expressed LIF, followed by IL-1α. Mice that had previously received LIF had increased peripheral PMNs and splenic HSPCs and GMPs upon IL-1α injection compared to empty vector controls (Fig. 7A-C). This data suggests that LIF exerts a functional impact on hematopoietic capacity that can potentiate the myelopoietic impact of IL-1α.

**Figure 7.**
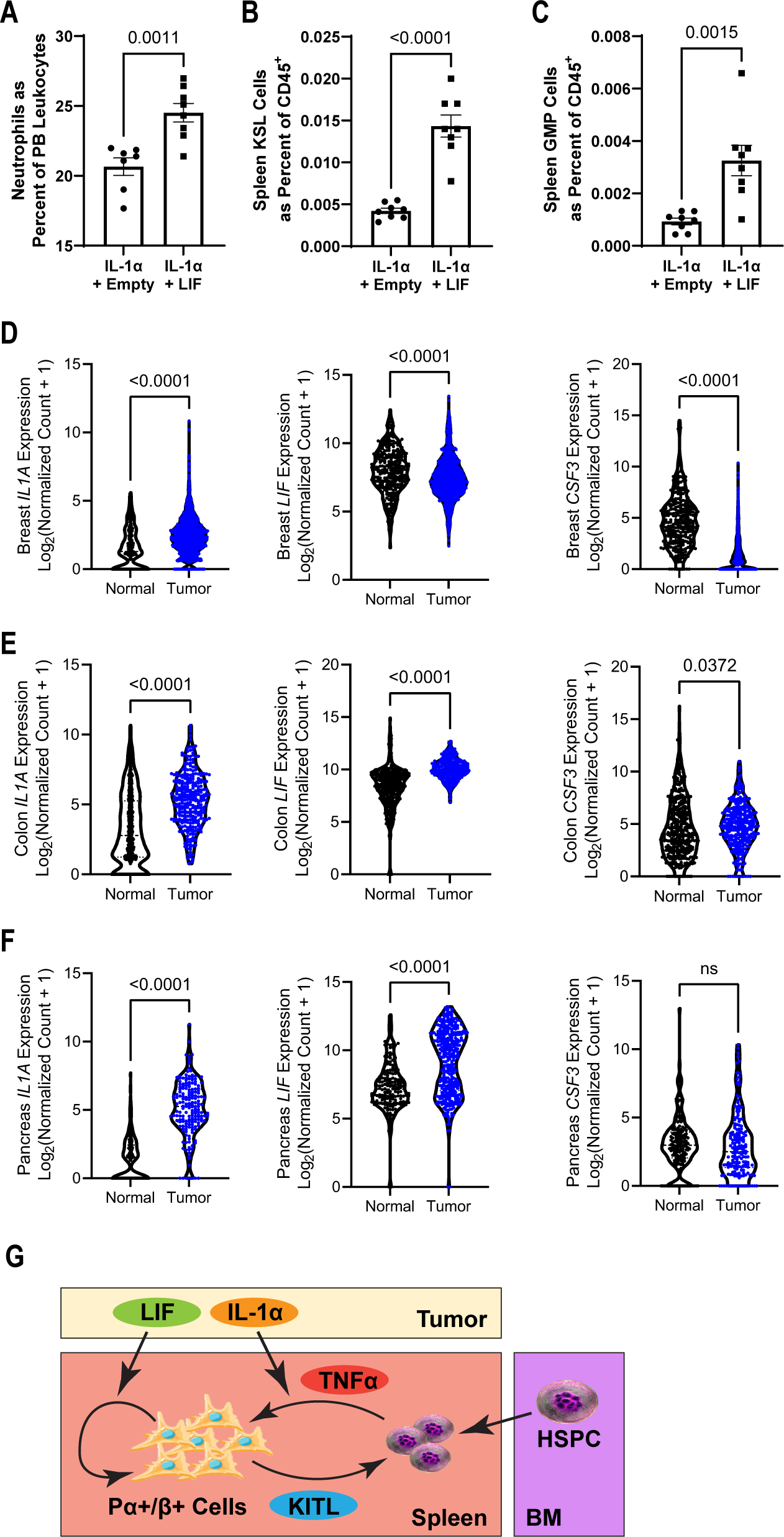
Human tumors co-express LIF and IL-1α which synergize in mouse models to potentiate EMH. (A – C) In mice with 10 days of LIF overexpression lentivirus or empty vector control lentivirus and 24 hours after treatment with 200ng IL- 1α i.v., PMNs in the PB as a percent of total leukocytes (A, n = 7-8), KSL cells as a fraction of total splenic CD45^+^ cells (B, n = 7-8), GMP cells as a fraction of total splenic CD45^+^ cells (C, n = 7-8). (D – F) RNA-seq expression of *IL1A*, *LIF*, and *CSF3* expression in tumor compared to normal tissue for breast (D, 292-1099), colonic (E, n = 288-349), and pancreatic (F, n = 171-178). (G) Our proposed model of parallel mechanisms for tumor associated EMH mediated in part by indirect inflammatory changes to HSPCs through IL-1α and direct proliferative effects on splenic ABS cells through LIF.

The PyMT-B6 mouse breast cancer line expresses IL-1α and LIF in addition to G-CSF. To examine whether the combination of IL-1α and LIF in the absence of G-CSF is relevant to human disease, we reexamined human tumor RNA-sequencing datasets from TCGA and other sources. We found that human breast cancer does not have the same cytokine profile as it only overexpresses IL-1α but not LIF or G-CSF (Fig. 7D, Fig. S4A), while human colon cancer was the only cancer to overexpresses all three cytokines (Fig. 7E). Importantly, human pancreatic, stomach, brain, and bile duct cancer all have overexpression of both IL-1α and LIF relative to normal tissue while having only minor changes in G-CSF (Fig. 7F, Fig. S5A-C). These data suggest that the co-occurrence of IL-1α and LIF is potentially clinically relevant for a diverse set of human cancers. Collectively, this illuminates a novel, potential mechanism by which human cancers may generate a myeloid-biased immune environment through EMH (Fig. 7G).

## Discussion

Extramedullary hematopoiesis (EMH) can be viewed as a process undertaken to meet the immense demand for myeloid cells during pathology that exceeds the capacity for existing bone marrow progenitors, making EMH a mechanism of emergency hematopoiesis. While EMH has been shown in a wide range of inflammatory conditions and diseases [3, 4, 8], the mechanisms regulating EMH have not been clearly elucidated. Here, we show the spleen as a critical site of EMH during solid tumor pathology that drives increases in peripheral blood neutrophilia. Consistent with this is recasting the spleen as a primary lymphoid organ involved in sensing systemic inflammation and activating by expanding total hematopoietic capacity, often with a myeloid bias. This framing of splenic function is concordant with data demonstrating an origin for myeloid cells within the spleen during various inflammatory and non- inflammatory pathologic states in both humans and mice [10, 11]. Previous work in a hepatocellular carcinoma mouse model has demonstrated that the absence of the spleen is sufficient to positively impact immune checkpoint blockade therapy [9].

Additional work has suggested that absence of the spleen leads to significantly fewer tumors developing in an inducible model of lung cancer [11]. We also show the reduction in the magnitude of tumor-induced neutrophil bias in the periphery following splenectomy. Interestingly, our study finds less profound alterations to the bone marrow compartment when compared to other investigations of solid tumor induced effects on hematopoiesis. In particular, Casbon *et al.* found a trend towards increased BM cellularity and progenitor numbers using the MMTV-driven PyMT transgenic breast cancer model [55]. Potentially, these discrepancies are the result of several differences between our experimentation and theirs including mouse genotypes, a transgenic versus tumor transplantation model, and a longer timeframe in the transgenic model.

Additionally, we also speculate that increases to BM HSPCs seen in the previously mentioned paper and others may contribute to increased splenic HSPCs through migration as observed in our model. These subtle differences and their impact on stem cell phenotypes highlight the nuance and limited understanding still present in the field of hematopoietic modulation by solid tumors.

While tumor manipulation of local immune cells within the tumor microenvironment has received significant attention, how tumor cells manage the immune system distally is less well-understood. In this paper, we demonstrate that profound expansion of hematopoiesis into the spleen occurs with breast cancer. We identify two cytokines produced by tumor cells that have distinct but overlapping interactions with splenic HSPCs and stromal cells to expand the size and functional capacity of the splenic niche to accommodate increased myelopoiesis. We present novel findings that support this conclusion. First, splenic HSPCs accompanying tumor presence express a gene profile characterized by TNFα. Second, IL-1α released by the tumor cells acts distally to induce TNFα expression in HSPCs. Third, tumors may indirectly activate splenic niche capacity in PDGFRα+/β+ stromal cells through local TNFα produced by inflammatory HSPCs.

Fourth, tumors directly expand the splenic niche through LIF by inducing proliferation in splenic PDGFRα+/β+ stromal cell populations. Moreover, LIF receptor deletion in PDGFRα+ cells significantly reduced hematopoietic capacity within the spleen. These data extend the role of this underappreciated cell type by centering it as the activatable niche cell within the spleen. Importantly, identifying LIF as expanding this cell type adds to our appreciation of stromal cells as active members of inflammatory pathology and supplements the roles LIF is already known to play in cancer. For instance, LIF is frequently overexpressed in many solid tumors including colorectal cancers, breast cancers and skin cancers and LIF overexpression in tumors correlates with poor prognosis of patients [56–58]. In mouse models, LIF blockade leads to reduced tumor progression [59–61]. Our analysis of human tumor data hints that LIF may also have local effects supporting cancer-associated fibroblasts, a cell-type which has recently drawn attention as key member of the tumor immune environment [62]. Collectively, we propose the parallel mechanisms of IL-1α and LIF that can synergize to activate splenic HSC niche to increase PMN production that may function in human cancers (Figure 7G).

Our data add depth and scope to the mechanisms by which cancers manipulate the host to generate a favorable immune environment for their growth, stretching as far up the differentiation hierarchy as primitive hematopoietic stem cells and their associated niche. One avenue that our paper focuses on is the cytokine axis established by tumor cells themselves. This focus uses an emerging classification of tumors by their functional effects that helps overcome heterogeneity both within and between tumor types and also makes comparisons of tumor pathology more congruous across species boundaries [63]. Studying tumor-derived cytokines also dovetails with the recent developments in understanding the reaction of HSPCs to inflammation [13, 64–67]. Our data also adds to the growing evidence supporting an active role in pathology played by HSPCs through inflammatory cytokine production [9, 68]. Many cases of cytokine-induced changes to HSPCs result in myeloid lineage bias. This shift towards the production of myeloid cells benefits tumor growth while tending to harm cancer patients. Across multiple tumor types, including breast, colon, pancreatic, and gastric cancer, as well as a systematic review of all cancer types, a high neutrophil-to-lymphocyte ratio is an independent prognostic factor for survival [17–20]. In addition to increased quantity, myeloid cells produced in communication with cancer cells have unique qualities that help drive cancer pathology, such as myeloid-derived suppressor cells [9]. Our data expand the function of inflammatory cytokines produced by HSPCs beyond myeloid lineage biasing. Particularly, we provide data showing that TNFα expressed by HSPCs can regulate the function of their own niche. In concert with tumor-produced LIF that expands the quantity of splenic HSPC niche cells, tumor-derived IL-1α induces TNFα expression by HSPCs to alter niche function into favoring increased EMH. This data warrants future studies addressing whether disruption of the local IL-1α/TNFα axis can impede EMH and how cell products of EMH induced by IL-1α and LIF impact the tumor microenvironment and cancer outcomes.

## Materials and methods

### Mice

Wild-type C57BL/6J mice (#000664), B6N.Cg-Tg(PDGFRa-cre/ERT)467Dbe/J (#018280), B6.SJL-Ptprca Pepcb/BoyJ (#002014), and B6.FVB-Tg(Cdh5-cre)7Mlia/J mice (#006137) were obtained from The Jackson Lab. MMTV-PyMT mice on a C57BL/6J background were a gift from Dr. M. Egeblad. *Lifr*-flox mice were obtained courtesy of Dr. Colin Steward [69]. All mice used in experimentation were female between the ages of 8 and 16 weeks unless otherwise stated. Animal husbandry, handling, and experimentation were approved by the Institutional Animal Care and Use Committee of Washington University School of Medicine.

### Mouse tumor models

MMTV-PyMT transgenic mice were used as a spontaneous model of breast cancer and were analyzed when evidence of peripheral neutrophilia was present which was between 3 to 6 months. For tumor transplantation studies, 5x10^5^ PyMT-B6 tumor cells, 5x10^5^ LLC tumor cells, 2x10^6^ 1956, or 2.5x10^5^ PyMT-B6 gene knockout tumors cells were injected subcutaneously in a slurry of 1:1 EHS ECM growth factor-reduced gel (Corning, # 354230; Sigma, #E6909) to PBS into the flank of the mouse and harvested after 21 days for PyMT-B6, 16 days for LLC, 17 days for 1956, and 28 days for PyMT- B6 gene knockout experiments. PyMT-B6, wild-type and knockout, cells and LLC cells were grown in DMEM with penicillin/streptomycin, 10% fetal bovine serum, and 10mM HEPES buffer. 1956 cells were grown in RPMI-1640 with penicillin/streptomycin, 10% fetal bovine serum, 100mM sodium pyruvate, 7.5% v/v sodium bicarbonate, and 50µM beta-mercaptoethanol. Supernatants were collected after 24 hours of incubation in culture starting 2 days after passage.

### Flow cytometry

Spleens were homogenized through a 100-μm filter. BM from femur and tibias was ejected by centrifugation at 3,200g for 2min at 4C. Peripheral blood was collected by cheek bleed. RBCs were lysed when needed using ACK lysis buffer (ThermoFisher, A10492-01). Cells were counted on an automated Nexcelom cell counter.

Cells were blocked with TruStain FcX PLUS anti-CD16/32 antibody (Biolegend, 156603) or anti-CD16/32 BV421 (Biolegend, clone 93) where appropriate before staining with antibodies followed by flow cytometry on a Gallios (Beckman Coulter) or a FACScan II (BD). When staining for intracellular cytokines, Cytofix/Cytoperm (BD, 554714) was used according to manufacturer’s instruction and 1μg/mL brefeldin A was maintained in the FACS buffer until fixation. Viability staining was added according to manufacturer’s instructions before beginning flow cytometry. Analysis was performed with FlowJo v10 software (Tree Star).

The following antibodies and reagents were purchased from BioLegend: anti-CD45.2 APC (clone 104), anti-CD11b APC-Cy7 (clone M1/70), anti-CD11b PE (clone M1/70), anti-Gr1 FITC (clone RB6-8C5), anti-Gr1 APC (clone RB6-8C5), anti-B220 PerCP/Cy5.5 (clone RA3-6B2), anti-B220 PerCP-Cy5.5 (clone RA3-6B2), anti-CD3e FITC (clone 145-2C11), anti-CD3e PE-Cy7 (clone 145-2C11), anti-Sca-1 APC (clone D7), anti-Sca-1 PerCP-Cy5.5 (clone D7), anti-CD45 AF700 (clone 30-F11), anti-CD45 BV421 (clone 30-F11), anti-c-Kit PE-Cy7 (clone 2B8), anti-c-Kit PE (clone 2B8), anti-VCAM-1 APC (clone 429), anti-PDGFRβ APC (clone APB5), anti-PDGFRα (clone APA5), 7-AAD dye (#420404), anti-IL-7R PE-Cy7 (clone A7R34), streptavidin PerCP- Cy5.5 (#405214), streptavidin BV421 (#405225), streptavidin APC (#405207), and biotin anti-lineage (#133307). Anti-CD34 FITC (clone RAM34) was purchased from Thermo.

Anti-CD45.1 PE (clone A20) was purchased from BD Biosciences. Anti-KITL biotin (#102501) and biotinylated goat IgG control (#105601) were purchased from R&D Systems.

Cell type delineations were made as follows: KSL cells were gated as CD45^+^/Lineage^-^/c-Kit^+^/Sca-1^+^; granulocyte-monocyte precursor (GMP) cells were gated as CD45^+^/Lineage^-^/c-Kit^+^/Sca-1^-^/CD16/32^+^/CD34^+^; common lymphoid progenitor (CLP) cells were gated as CD45^+^/Lineage^-^/c-Kit^-^/Sca-1^+^/IL-7R^+^.

### Colony forming assay

Peripheral blood or the full contents of ABS:hematopoietic progenitor co-culture 24- wells were plated into complete methylcellulose media (Stem Cell Technologies, M3434). Colonies were scored 7-14 days after plating.

### Bone marrow transplant

For splenocyte transplantation, CD45.2 mice were irradiated with 9.5 Gy and 1x10^6^ splenocytes from CD45.1 control or CD45.1 tumor-bearing animals were injected intravenously by the retroorbital route 24 hours after irradiation. For niche function studies, CD45.2 mice were irradiated with 9.5 Gy and CD45.1 hematopoietic cells were isolated from cell culture with or without ABS cells and injected intravenously by the retroorbital route 24 hours after irradiation. Mice were monitored daily for mortality or signs of severe morbidity up to 28 days. Mice were maintained until mortality to evaluate the long-term reconstitution potential.

### Splenectomy

Splenectomies and sham surgeries were conducted courtesy of the Hope Center Animal Surgery Core, Washington University School of Medicine. After a week recovery period, mice were injected with PyMT-B6 tumor cells as detailed above.

### Single cell RNA-sequencing and analysis

Spleens were minced and digested in 1mg/mL Collagenase Type IV + 0.25mg/mL DNase I. Bone marrow was removed by centrifugation as detailed above and digested. Digestion was quenched then filtered through a 100μm filter. Cells were pelleted, counted, and aliquoted. TruStain FcX™ PLUS was used to block samples then biotin anti-lineage antibodies were used to stain lineage cells. After washing, strepavidin magnetic beads (NEB, S1420S) were used to deplete lineage positive cells. Remaining cells were pelleted and then stained with streptavidin BV605 (Biolegend, #405229), anti- CD45 AF700, anti-PDGFRα APC, anti-CD51 PE (Biolegend, clone RMV-7), anti-CD31 PE-Cy7 (Biolegend, clone 390), anti-Sca-1 PerCP-Cy5.5 (Biolegend, clone D7), and anti-c-Kit FITC (Biolegend, clone 2B8). Cells were then washed into holding buffer (0.04% BSA in PBS), stained with DAPI, and sorted on a high modified MoFlo into five populations: Live/Lin^-^/CD45^+^/c-Kit^+^/Sca-1^+^, Live/Lin^-^/CD45^+^/c-Kit^+^/Sca-1^-^, Live/Lin^-^/CD45^-^/CD31^+^, Live/Lin^-^/CD45^-^/CD31^-^, Live/Lin^-^/CD45^-^/CD31^-^/CD51^+^, Live/Lin^-^/CD45^-^ /CD31^-^/CD51^-^. These populations were combined at equal ratios and submitted for 10X Genomics 3’ v3.1 Chemistry sample preparation and sequencing on a NovaSeq6000 at the Genome Technology Access Center.

Cell Ranger (10x Genomics, Pleasanton, CA) with default settings de-multiplexed, aligned, filtered, and counted barcodes and UMIs. SoupX preprocessing was used to remove ambient RNA contamination at a contamination fraction of 10% [70]. Filtered outputs were imported into R v4.0.5 using Seurat v3.2.3 and barcodes with fewer than 350 unique genes were excluded. Seurat objects from the four experiment groups were merged and an SCT transformation with a variable feature count of 20,000 was performed on the resulting object.[71, 72] The dimensions of the object were reduced using RunPCA with principal coordinates equal to 50. UMAP coordinates were calculated using all 50 PCA dimensions and a minimum distance of 0.05.

FindNeighbors function was used to compute nearest neighbors using all 50 PCA dimensions and FindClusters function at a resolution of 1.2 was used to compute cell clusters. Markers for each cluster were calculated using FindAllMarkers function with a minimum percentage of 0.1.

For reanalysis of a publically available single cell RNA-sequencing dataset of bone marrow niche cells [47], data was downloaded from GSE108891 on Gene Expression Omnibus. Raw counts files for GSM2915575, GSM2915576, GSM2915577, and GSM3330917 were imported into R using Seurat 3.2.3 and barcodes with fewer than 500 unique genes were excluded. Seurat objects from the four experiment groups were merged and an SCT transformation with a variable feature count of 8,000 was performed on the resulting object [71, 72]. The dimensions of the object were reduced using RunPCA with principal coordinates equal to 20. UMAP coordinates were calculated using all 20 PCA dimensions and a minimum distance of 0.05.

FindNeighbors function was used to compute nearest neighbors using all 20 PCA dimensions and FindClusters function at a resolution of 0.2 was used to compute cell clusters. Markers for each cluster were calculated using FindAllMarkers function on default settings.

### Magnetic bead isolation and quantitative reverse transcriptase analysis

TruStain FcX™ PLUS antibody was used to block samples then biotin anti-lineage antibodes and biotin anti-Flk1 (Biolegend, clone 89B3A5) antibody were used to stain cells. After washing, strepavidin magnetic beads were used to bind the stained cells. Positive cells were depleted by two rounds of magnetic selection. Depleted cells were pelleted and stained with anti-CD34 FITC and anti-FITC biotin (Biolegend, clone FIT-22). Cells were washed, pelleted, and resuspended before adding streptavidin magnetic beads. After incubation, the tubes were placed on the magnet and the supernatant removed. Using an RNeasy Kit Micro (Qiagen, #74004), RLT buffer was used to lyse the cells before proceeding with RNA isolation according to manufacturer’s instructions. qScript™ cDNA SuperMix (QuantaBio, 95048-100) was used to produce cDNA before running RT-qPCR with 2x SYBR Green qPCR Master Mix (BiMake, B21203) according to manufacturer instructions. Primers sequences were as follows: *Tnf* forward - CCCTCACACTCAGATCATCTTCT, reverse - GCTACGACGTGGGCTACAG; *Cxcl2* forward – CCAACCACCAGGCTACAGG, reverse – GCGTCACACTCAAGCTCTG;*Nfkbia* forward – TGAAGGACGAGGAGTACGAGC, reverse –TTCGTGGATGATTGCCAAGTG; *Nfkbiz* forward – GCTCCGACTCCTCCGATTTC, reverse – GAGTTCTTCACGCGAACACC; *Mki67* forward – ATCATTGACCGCTCCTTTAGGT, reverse – GCTCGCCTTGATGGTTCCT; *Il1r1*forward – GTGCTACTGGGGCTCATTTGT, reverse – GGAGTAAGAGGACACTTGCGAAT; *Hprt* forward – TCAGTCAACGGGGGACATAAA, reverse – GGGGCTGTACTGCTTAACCAG.

### ELISA and multiplex protein assay

ELISA kits for IL-1α (Abcam, ab199076), CXCL1 (R&D, DY453-05), and CXCL12 (Abcam, ab100741) were used according to manufacturers’ instructions. LIF serum samples were analyzed using the Abcam, ab238261, while all other sample types were analyzed using R&D, DY449. Serum samples from MMTV-PyMT mice and littermate controls were sent to Eve Technologies (Calgary, AB, Canada) and assayed using the 44-plex Mouse Discovery assay. Results from Eve Technologies were imported into R, log10 normalized, and plotted using the heatmap.2 function in the gplots package.

### In-vivo cytokine injection

TNFα (Peprotech, 315-01A) and IL-1α (Peprotech, 211-11A) was purchased, resuspended according to manufacturer’s instructions. For TNFα and IL-1α experiment, 2μg and 0.5μg or 0.2μg per mouse were injected retroorbitally, respectively. Mice were analyzed 24 hours later.

### CRISPR-Cas9 gene deletion in PyMT-B6 Cells

PyMT-B6 cells were seeded and then grown overnight to around 70% confluence before adding TrueCut™ Cas9 Protein v2 (Thermo, A36497), Lipofectamine™ CRISPRMAX™ Cas9 Transfection Reagent, and TrueGuide™ Synthetic sgRNA (Thermo, #A35533) according to manufacturer’s instructions. Guide RNAs from the manufacturers catalog were selected to be positioned in the earliest exon shared by all known isoforms and to minimize the distance between the two cut sites. Both guides were incubated with the cells during lipofection. After lipofection, cells with single cell cloned. Each clone was tested for deletion of the gene by ELISA, Sanger sequencing, and gel electrophoresis when applicable. *Csf3* was deleted initially then a successful clone was used as the parental line for subsequent deletion of *Lif* or *Il1a*. These knockout cell lines were injected *in vivo* as described above.

### Splenic stromal cell isolation, culture, and co-culture with hematopoietic progenitors

Spleens were minced and plated on gelatin coated plates. Growth media for cells was alpha-MEM with 10% FBS, 1x Glutamax, 10mM HEPES buffer, 100μg/mL Primocin (InVivogen, ant-pm), and 5ng/mL heat stable FGF2 (Gibco, PHG0368). After 72 hours, non-adherent tissue was gently removed. Media was changed every 2-3 days thereafter until the culture was 100% confluent. Cells were passaged using CellStripper and plated without gelatin coating. For flow cytometry experiments involving membrane KITL staining, cells were lifted using CellStripper and stained. For other flow cytometry experiments and cytokine stimulation, cells were lifted with Trypsin-EDTA. For LIF stimulation experiments, cells were plated at 5,000 cells/cm^2^, grown overnight in growth media, then changed to growth media without heat-stable FGF2 with or without 20ng/mL LIF (Peprotech, 250-02). Media was changed after two days and the RNA was harvested on the third day. For TNFα stimulation experiments, cells were plated at 10,000 cell per cm^2^, grown over night in growth media, then changed to growth media with or without 2.5ng/mL of TNFα (Peprotech, 315-01A). Cells or supernatant were harvested after 24 hours for flow cytometry or ELISA, respectively.

For co-culture with hematopoietic stem and precursor cells, splenic stromal cells were plated and grown until confluence before 5,000 live c-Kit^+^ Lineage^-^ cells were sorted and transferred into individual 24-wells with or without a stromal monolayer. Co-cultures were then grown for 7 days before passage or usage in an experiment as specified. The same media was used for co-culturing as was used for monoculture of splenic stromal cells.

### Lentiviral particle production and administration

Murine LIF ORF (NM_008501.2) was purchased from GenScript and cloned into the pCSII-EF1α-IRES2-bsr lentiviral backbone. Lentiviral packaging plasmid psPAX2 (Addgene, plasmid #12260) and VSV-G envelope expressing plasmid PMD2.G (Addgene, plasmid #12259) were gifts from Didier Trono. 293FT cells were transfected with lentiviral DNA using the calcium phosphate method. Virus was concentrated from media using PEG Virus Precipitation Kit (Sigma). Viral titer was determined by QuickTiter™ Lentivirus Associated HIV p24 Titer Kit (Cell Biolabs, INC). Mice were infected by tail vein injection with 4x10^9^ viral particles before sacrifice on day 7 for immunofluorescence experiments or on day 10 for all other experiments.

### Immunofluorescence, bright-field, and confocal microscopy

For immunofluorescence, spleens were removed from animals and directly embedded by freezing into NEG-50 media. 6μm sections were fixed using 4% PFA in PBS then permeabilized in 0.5% Triton-X100 in PBS before blocking with 1% BSA/ 5% donkey serum in PBS. Sections were stained with primary antibodies overnight and then stained with secondary antibodies for 1 hour. Primary antibodies, anti-PDGFRa (AB Online, # ABIN726620) and anti-Ki67 (Biolegend, clone 16A8), were diluted 1:200 for staining. Secondary antibodies were donkey anti-rabbit AF488+ (ThermoFisher, # A32790) and donkey anti-rat AF594+ (ThermoFisher, #A21209). Sections were quenched using ReadyProbes™ Tissue Autofluorescence Quenching Kit (ThermoFisher, R37630) according to manufacturers’ instructions before staining with DAPI and mounting with ProLong™ Diamond Antifade Mountant (ThermoFisher, P36970). Slides were sealed and imaged using a Zeiss AxioImager Z2 at the Washington University Center for Cellular Imaging using Zen Blue v.3 for image acquisition and processing. Images were counted manually.

For bright-field microscopy, day 7 stromal:hematopoietic co-cultures were imaged live on an ACCU-SCOPE EXI-600 inverted microscope. Images were processed using ImageJ [73].

For confocal microscopy, spleens were removed from animals and fixed in 4% PFA (Electron Microscope Sciences, #15710-S) with PBS for 72hrs. Spleens were washed overnight in PBS and then sectioned by Vibratome to 300μm. Sections were then cleared using 10% w/v CHAPS and 25% v/v N-Methyldiethanoamine in PBS for 48hrs before washing with PBS followed by 72hrs of blocking 5% donkey serum (Sigma, #D9663) in PBS. Primary antibodies, anti-PDGFRa (AB Online, # ABIN726620), anti-Kitl (R&D, #AB- 455-NA) and anti-c-Kit (Biolegend, clone 2B8) were then stained at a 1:200 dilution for 72hrs. Sections were washed with PBS overnight before staining at 1:250 with secondary antibodies, donkey anti-rat AF647+ (Thermo, # A48272), donkey anti-rabbit AF555 (Thermo, A-31572), and donkey anti-Goat AF405+ (Thermo, # A48259). After secondary staining, sections were washed overnight with PBS before dehydration with increasing concentrations of ethanol – 50%, 70%, 95%, and 95% - for at least 2 hours each before incubation with a methyl salicylate solution (Sigma-Aldrich, M6752) for 30-60 minutes in a custom metal chamber with 0.2mm coverslip glass bottom. Tissue sections were then imaged at 1.5µm optical sections using a seven-laser inverted Leica SP8 microscope with full spectral hybrid detectors. All image collection was performed using Leica LAS X software, and analysis was performed using Leica LAS X or Imaris (Bitplane) v8 and v9 software. Images shown are maximum intensity projections of 8 sections representing 10.5µm in depth.

### Human tissue datasets and xCell analysis

Transcriptomic data of tumor and normal samples were downloaded from The Cancer Genome Atlas (TCGA), Therapeutically Applicable Research To Generate Effective Treatments (TARGET), and Genotype-Tissue Expression (GTEx) consortiums were downloaded using the UCSC Xena portal (https://xena.ucsc.edu/). Normalized RSEM expected counts were logged for visualization and statistical purposes.

A signature based deconvolution pipeline, xCell [74], was used to identify enrichment of stromal populations in the tumor microenvironment. Gene length normalized TPM data from TCGA was downloaded from the UCSC Xena portal was used as an input into xCell for stromal cell deconvolution. Patients were grouped into quartiles by LIF expression and compared across subgroups.

### Quantification and statistical analysis

Statistical analyses were performed using GraphPad Prism 9 software (GraphPad Software, San Diego, CA). P values were calculated using unpaired t tests (two-tailed) unless otherwise indicated in the figure legends. P values less than 0.05 was considered statistically significant and displayed above the comparison bars in figures. Each figure represents at least two independent experiments and are presented together unless otherwise specified. Error bars show the standard error of the mean for each sample.

### Data sharing statement

The single cell RNA-sequencing data reported in this paper will be uploaded to GEO, and the accession number will be given before time of publication. All data from raw sequencing reads to analyzed data along with the accompanying code where applicable is available to reviewers upon request.

## Authorship contributions

D.A.G.B. and K.C. conceived the study, analyzed the data, and wrote the paper. D.A.G.B. and J.W. performed the *in vivo* experiments. D.A.G.B. analyzed the scRNA- seq experiments. D.A.G.B., M.K., and A.K. conducted *in vitro* experiments. D.A.G.B. and M.S. analyzed human RNA-seq data. D.A.G.B. and K.K. created and purified lentivirus. D.A.G.B. and B.Z. imaged the spleen by confocal microscopy. C.L.S. provided the *Lifr*-flox mice.

## Acknowledgements

We thank Drs. M. Egeblad and David DeNardo for the gift of the MMTV-PyMT-B6 mice and PyMT-B6 cell line, respectively. We thank Dr. Gwendalyn Randolph for the use of her instruments and laboratory space for confocal imaging. We also thank the TCGA Research Network for providing data used in this publication. This work was supported by the NIH T32 AI007163 (D.A.G.B), the NIH grants R37AI049653 (B.H.Z.), R01HL149954, R01HL55337, and Siteman Investment Program Research Development Awards (K.C.).

## Abbreviations

HSPC: hematopoietic stem and precursor cell
EMH: extramedullary hematopoiesis
LIF: leukemia inhibitory factor
BM: bone marrow
PMN: polymorphonuclear leukocytes neutrophils
PB: peripheral blood
KSL: c-Kit^+^Sca-1^+^Lineage^-^ cells
GMP: granulocyte-monocyte progenitor
CLP: common lymphoid progenitor
MMTV: murine mammary tumor virus
PyMT: polyomavirus middle T antigen.

